# Brain structural differences between 73- and 92-year olds matched for childhood intelligence, social background, and intracranial volume

**DOI:** 10.1101/135871

**Authors:** Stuart J. Ritchie, David Alexander Dickie, Simon R. Cox, Maria del C. Valdés Hernández, Alison Pattie, Devasuda Anblagan, Paul Redmond, Natalie A. Royle, Janie Corley, Susana Muñoz Maniega, Adele M. Taylor, Sherif Karama, Tom Booth, Alan J. Gow, John M. Starr, Mark E. Bastin, Joanna M. Wardlaw, Ian J. Deary

## Abstract

Fully characterizing age differences in the brain is a key task for combatting ageing-related cognitive decline. Using propensity score matching on two independent, narrow-age cohorts, we used data on childhood cognitive ability, socioeconomic background, and intracranial volume to match participants at mean age 92 years (*n* = 42) to very similar participants at mean age 73 (*n* = 126). Examining a variety of global and regional structural neuroimaging variables, there were large differences in grey and white matter volumes, cortical surface area, cortical thickness, and white matter hyperintensity volume and spatial extent. In a mediation analysis, the total volume of white matter hyperintensities and total cortical surface area jointly mediated 24.9% of the relation between age and general cognitive ability (tissue volumes and cortical thickness were not significant mediators in this analysis). These findings provide an unusual and valuable perspective on neurostructural ageing, in which brains from the eighth and tenth decades of life differ widely despite the same cognitive, socio-economic, and brain-volumetric starting points.

## Introduction

Many changes in brain structure occur during normal ageing. Understanding and characterizing these age-related differences is important, since they have been linked to ageing-related cognitive decline, a pervasive and growing phenomenon with a substantial predicted effect on ageing societies (Brayne, 2007; Luengo-Fernandez, Leal, & Gray, 2010). Relatively few studies have modeled both brain and cognitive age differences, and fewer have included participants over the age of 90 years (Dickie et al., 2013). In the present study, we quantify age differences in a variety of neurostructural measures using an unusual design; we compare closely-matched participants from two independent narrow-aged samples in later life, one aged 73 and the other aged 92 years. We then test the extent to which neuroanatomical differences can explain the large age-related cognitive differences between the two samples.

The most-studied neuroanatomical measure with reference to ageing is brain volume. Volume peaks in early adulthood, before a period of relatively mild decline through midlife and more rapid degeneration in older age (Fjell & Walhovd, 2010). In non-pathological ageing, adults aged over 60 experience around 0.5% decline in total brain volume per year (Fotenos et al., 2005), with volumetric declines seen in both grey and white matter, in regions across the entire brain (Dickie et al., 2015; Giorgio et al., 2010; Kruggel, 2006; Raz et al., 2005; Walhovd et al., 2011; Ziegler et al., 2011). Cortical surface area follows a similar trajectory of decline (Hogstrom et al., 2013), and most regions of the brain exhibit cortical thinning with age, with the loss of up to ∼0.6mm of cortical thickness per decade (Thambisetty et al. 2010; see also Fjell et al., 2009; Shaw et al., 2016a,b). Finally, the volume of white matter hyperintensities (WMH) tends to increase with advancing age (Morris et al., 2009; Wardlaw et al., 2015; Ritchie et al., 2015a).

Deteriorations in the above-listed brain measures have been linked, in longitudinal studies, to declines in key cognitive faculties such as fluid intelligence, reasoning, mental speed, and memory, which decline on average throughout adulthood (Salthouse, 2004). For example, Schmidt et al. (2005) showed that declining brain volume was related to loss of cognitive skills such as memory and visuopractical abilities (see also Jokinen et al., 2012; Ritchie et al., 2015a). In a meta-analysis, Kloppenborg et al. (2014) showed that advancing WMH levels were related to decrements in all measured cognitive abilities.

There is relatively little evidence on which of these neuroanatomical variables are the most relevant for explaining cognitive ageing: few studies have analysed multiple imaging variables simultaneously. In a previous study of one of the cohorts involved in the present analysis (Ritchie et al., 2015b), we measured multiple neuroanatomical measures and related them to a broad latent variable of general cognitive ability (so-called ‘*g*’) measured at age 73 years. Total brain volume made the largest contributions to explaining variance in *g*, but other variables such as WMH and cortical thickness made additional, incremental contributions (see also Kievit et al., 2012). Thus, it is likely that several different aspects of brain structure are independently relevant to understanding the ageing of cognitive abilities. However, these studies focus on cognitive ability level, rather than the age-related differences in these abilities.

In testing the extent to which brain structure can account for age differences in cognitive functioning, the present study takes the approach of Kievit et al. (2014). They used structural equation model-based mediation analysis to test whether the age variance in cognitive ability could, in part, be explained by different neuroanatomical measures. They showed, for instance, that fractional anisotropy of the forceps minor and the volume of Brodmann Area 10 were parallel mediators (explaining 18.2% in total) of the association between age and fluid intelligence in a sample with an age range of 18 to 89 years. Though the selection of brain regions included in that analysis were limited (two cortical regions and two tracts), their results contributed to our understanding of the multifaceted nature of brain ageing and its relation to key cognitive outcomes. That, in addition to a detailed characterization of ageing across various brain imaging measures, is the aim of the present study.

### The Present Study

Here, we extensively characterize whole and regional brain differences between two narrow-age cohorts of older people, one aged around 73 years and the other aged around 92 years. Unusually, both cohorts had data available on the same well-validated general cognitive ability test taken at age 11 years, as well as retrospective data on their socioeconomic status from childhood and adulthood. We used propensity score matching on these background variables, as well as on a measure of maximal brain size (their intracranial volume), to reduce confounding in the comparison of the two cohorts in later life.

Using these well-matched cohorts, we ran the following three analyses. First, we characterized the extent of the 19-year age differences in multiple broad brain volumetric measures: total brain volume, grey and white matter volume, and the volume of white matter hyperintensities. Second, we examined grey matter in more detail, using parcellation to map volume and surface area differences in each of 54 grey matter regions of interest between the 73- and 92-years-olds. We also used a vertex-wise method to examine the cohort differences in cortical thickness across the entire brain. Third, we used mediation analyses to test whether differences in *g* (indicated by the same three cognitive tests taken by both cohorts) between the samples could be accounted for by differences in brain structure.

## Method

### Participants

Members of both the Lothian Birth Cohort 1921 (LBC1921; Deary et al., 2004) and the Lothian Birth Cohort 1936 (LBC1936; Deary et al., 2007, 2012) studies were included in the present analysis. Both cohorts are studies of ageing that follow individuals who, at age 11 years, took part in the Scottish Mental Surveys of 1932 or 1947. The cohorts have been followed-up at multiple waves in later life; for the present study, we focus on data from the fifth wave of the LBC1921 (total *n* = 59, mean age = 92.1 years, SD = 0.34) and the second wave of the LBC1936 (total *n* = 866, mean age = 72.5 years, SD = 0.71). At these waves, *n* = 53 members of the LBC1921, and *n* = 731 members of the LBC1936, attended for a structural MRI scan (as described below, the final matched sample involved *n* = 42 LBC1921 members and *n* = 126 LBC1936 members). In the LBC1921 cohort, cognitive/medical testing and brain scanning were completed on the same day in all but a few cases (mean gap = 0.04 days, SD = 0.27). In the LBC1936, the participants all made two separate visits (mean gap = 65.04 days, SD = 39.57).

Approval for the LBC1921 study was obtained from the Lothian Research Ethics Committee (Wave 1: LREC/1998/4/183; Wave 3:1702/98/4/183) and the Scotland A Research Ethics Committee (waves 4 and 5:10/S1103/6). Approval for the LBC1936 study was obtained from the Multi-Centre Research Ethics Committee for Scotland (wave 1: MREC/01/0/56), the Lothian Research Ethics Committee (wave 1: LREC/2003/2/29), and the Scotland A Research Ethics Committee (waves 2 and 3: 07/MRE00/58). All participants provided written, informed consent before any measurements were taken.

### Measures

#### Brain MRI acquisition and volumetric processing

Brain MRI acquisition parameters were described in detail for the LBC1936 by Wardlaw et al. (2011). All subjects (from both cohorts) had brain MRI on the same 1.5 Tesla GE Signa Horizon HDx clinical scanner (General Electric, Milwaukee, WI, USA), maintained on a careful quality assurance programme. Both cohorts underwent the same structural imaging examination consisting of high resolution 3D T1- weighted volume, and T2-, T2*-, and fluid attenuated inversion recovery (FLAIR)-weighted sequences. The voxel resolution of the T2-, T2*- and FLAIR-weighted MRI sequences was 2, 2 and 4 mm^3^ for the LBC1936, and 2.2, 2.2 and 4.4 mm^3^ for the LBC1921; the voxel resolution for the 3D T1-weighted volume scan (1.3 mm^3^) was identical between cohorts.

We measured intracranial, whole brain, grey matter, normal appearing white matter, and white matter hyperintensity (WMH) volumes in cm^3^ using a validated multispectral image processing method that combines T1-, T2-, T2*-, and FLAIR-weighted MRI sequences for segmentation (Valdés Hernández, et al., 2010). All sequences were co-registered and tissue volumes estimated by cluster analysis of voxel intensities. We explicitly defined WMH as punctate, focal or diffuse lesions in all subcortical regions and manually edited WMH masks according to STandards for ReportIng Vascular changes on nEuroimaging (STRIVE) guidelines (Wardlaw, et al., 2013). Editing was overseen by a consultant neuroradiologist (author J.M.W.). We manually checked all segmented images for accuracy blinded to all clinical details, corrected errors, and excluded imaging-detected infarcts from WMH volumes (Wang, et al., 2012). We tested the correlation of the overall grey matter volume produced using this method with the overall grey matter estimate derived using Freesurfer, which we used for our cortical subregional analysis (see below). Across the participants included in the present study, this correlation was *r*(164) = .89.

#### Lesion distribution maps

To produce a map of the distribution of WMH across the brain (as per Valdés Hernández et al., 2015), we co-registered all WMH masks to a common normal-ageing brain template (Dickie et al., 2016a) using affine registration on FSL-FLIRT (Jenkinson et al., 2002), averaged all the co-registered WMH masks of the datasets from each cohort (i.e. from LBC 1921 and LBC1936) and generated two probability distribution maps of WMH. Then, we mapped both probability maps in the template to analyze the spatial distribution of WMH from each sample. Illustrations of the distributions were produced in Mango v4.0 (http://ric.uthscsa.edu/mango/).

#### Subregional volumes and surface areas

FreeSurfer v5.3 (http://surfer.nmr.mgh.harvad.edu/) was used for cortical reconstruction and segmentation of T1-weighted volumes according to the Desikan-Killiany parcellation protocol (Desikan et al., 2006). Briefly, the steps involved removal of non-brain tissue, intensity normalization, tessellation of the white/grey matter boundary, inflation and registration of the cortical surface to the spherical atlas according to the folding patterns of each individual (Segonne et al., 2004; Segonne et al., 2007; Fischl and Dale, 2000; Fischl et al., 2004). This yielded the volume and surface area of 54 cortical regions of interest, where the surface area represents the sum of all triangular tessellation in each anatomical regions at the mid-point between grey and white matter. All output was visually inspected for processing failures or deficiencies (including tissue identification and boundary positioning errors), which failed in 3 cases in the LBC1921 (leaving *n* = 41 for the volume and surface area comparisons) and 4 cases in the LBC1936 (leaving *n* = 122). Cortical surface visualisations used the freely-available Liewald-Cox Heatmapper tool (http://www.ccace.ed.ac.uk).

#### Cortical thickness measurement

Cortical thickness was measured using the fully automated CIVET v1.1.12 image processing pipeline developed at the Montreal Neurological Institute (Ad-Dab’bagh, et al., 2006, Zijdenbos, et al., 2002). CIVET measures cortical thickness at 81,924 vertices (the perpendicular distance between grey and white matter surfaces) across the cortex (Ducharme, et al., 2016, Karama, et al., 2009, 2015). For clarity, we use ‘vertex’ to refer to the perpendicular distance between grey and white matter surfaces, not the cranial vertex. Cortical maps from each subject were manually inspected according to previously validated standards (Ducharme, et al., 2016). Approximately 10% of subjects from each cohort failed cortical thickness processing due to poor scan quality or motion artefacts. None of the failures in LBC1921 were part of the final *n* = 42 matched sample, but 14 subjects from the LBC1936 matched sample failed processing (thus, *n* = 112 were used in the cortical thickness analysis).

Cortical vertex-wise regression analyses were performed using the SurfStat MATLAB toolbox (http://www.math.mcgill.ca/keith/surfstat). The statistical significance of results for cortical thickness were corrected for multiple comparisons using Random Field Theory (RFT) to avoid false positives when more than 80,000 tests were performed (Chung et al., 2005). RFT identifies statistically significant “clusters” of vertices and vertex “peaks”. Cluster *p*-values show regions of connected vertices with *p*-values below .001 in clusters whose extent is significant at *p* < .05 (http://www.math.mcgill.ca.keith/surfstat), i.e., a collection of connected vertices with *p* < .001 that was unlikely to occur by chance. Vertex *p*-values show individual vertices where individual *t* scores are above the vertex-wise RFT critical *t*-value which is derived via the expected Euler characteristic (EC ≈ critical *p* value [0.05]) and number of resolution elements (“resels”) in the *t* cortical map (Brett et al., 2004; Worsley et al., 2004).

#### Matching variables

We ensured the cohorts were comparable by matching them on five variables that indexed their social, cognitive, and neural background (the propensity score matching procedure is described in the ‘Statistical Analysis’ section below). First, as noted above, all participants in both cohorts had childhood cognitive testing data available from age 11 years based on the same test. The test used, the Moray House Test No. 12 (Scottish Council for Research in Education, 1933), assesses a variety of cognitive functions with an emphasis on verbal reasoning, and is strongly correlated with other commonly-used tests of general cognitive function in childhood and later life (Deary et al., 2004). Second, each participant’s father’s socioeconomic status when the participant was aged 11 was collected by questionnaire around the time of the first wave of each study (at about 79 years for LBC1921 participants, and 70 years for LBC1936 participants), and classified using an index of their occupational class on a scale of I (professional) to V (unskilled; General Register Office, 1956). For female cohort members, the highest occupational class of the household was taken. Third, information on the participants’ own socioeconomic status was collected during the same interview; it was then classified, based on their most prestigious occupation they held before retirement, on the same scale for the LBC1921 participants and a similar scale for the LBC1936 (Office of Population Censuses and Surveys, 1980). Fourth, we matched the participants for intracranial volume, a proxy measure of maximal lifetime brain size. Intracranial volume, unlike total brain tissue volume (which shows steep declines across adulthood), has been shown to be highly stable between ages 18 and 96 (see Figure 2 in Royle et al., 2013). Its measurement in the present samples is described above. Fifth, we matched the participants for sex, such that the sex ratio within the matched sample of each cohort was similar.

#### Later-life cognitive tests

Members of both cohorts completed multiple cognitive tests in later life. For the purposes of this study, we took three tests that were administered in exactly the same setting in both cohorts and that could be expected to decline with age (that is, they measured ‘fluid’-type aspects of cognitive ability; see Salthouse et al., 2004). The first test was Digit-Symbol Substitution from the Wechsler Adult Intelligence Scale, 3^rd^ UK Edition (WAIS-III^UK^; Wechsler, 1998a), a measure of cognitive processing speed. The second was the Logical Memory test (Story A only) from the Wechsler Memory Scale-Revised in the LBC1921 (WMS-R; Wechsler, 1987) and the Wechsler Memory Scale, 3^rd^ UK Edition in the LBC1936 (WMS-III^UK^; Wechsler, 1998b), a test of verbal declarative memory. The third was a phonemic Verbal Fluency Test (using the letters C, F, and L, for one minute each; Lezak, 2004), which taps an aspect of executive function. We used structural equation modeling (see below) to estimate a general (*g*) factor of cognitive ability from these three tests.

Each participant was also administered the Mini-Mental State Examination (MMSE; Folstein, Folstein, & McHugh, 1975), a dementia screening instrument that assesses basic aspects of language, attention, and orientation to place and time, with a maximum score of 30. Scores below 24 are commonly considered indicative of possible mild cognitive impairment or dementia.

#### Health variables

A variety of self-reported (categorical) and blood-derived (continuous) health variables were assessed identically in both cohorts. The self-reported variables included smoking status (never, ex-, or current), self-rated health both at present (from poor to excellent) and compared to one year ago (much better to much worse), and diagnoses of a variety of conditions including hypertension, diabetes, cardiovascular disease, and stroke. The blood-derived biomarkers consisted of blood haemoglobin, white cell count, fibrinogen, D-dimer, estimated glomerular filtration rate, and glycated haemoglobin.

### Statistical analysis

#### Propensity score matching

The main aim of the study was to compare brains of young-old and old-old people who had similar cognitive, social, and brain-volumetric starting points. Participants were matched across the two cohorts using the five matching variables described above (childhood Moray House Test cognitive score, father’s social class, the participant’s own achieved social class, intracranial volume, and sex). To do this, we used propensity score matching in the ‘nonrandom’ package for R (Stampf, 2014). There were 42 LBC1921 participants who had full data on all the matching variables and who had attended for an MRI scan; for each of these, we selected three matching LBC1936 participants who also had all matching variables available and attended scanning (thus 126 LBC1936 participants). Participants were matched if they were within 0.2 standard deviations of the logit of the propensity score produced from a simultaneous logistic regression model including all five matching variables. We ensured that the matching was adequate by comparing across the samples on each of the matching variables using Welch’s two-sample *t*-tests, calculating Cohen’s *d*_*s*_ for the standardized effect size, as described by Lakens (2013). The same procedure was used to compare the samples on each of the brain variables of interest.

#### Mediation analysis

Another aim of the study was to test the extent to which general fluid cognitive differences between young-old and old-old people are accounted for by brain structural differences. We used structural equation modeling in the “lavaan” package for R (Rosseel, 2012) to perform mediation analysis. The predictor variable was the cohort to which each participant belonged (i.e. a proxy for whether they were 73 or 92 years old), the outcome variable was the *g*-factor of cognitive ability estimated from the three cognitive tests, and the potential mediator variables were the brain measures.

First, we tested whether the *g*-factors within each cohort were comparable, by testing their cross-cohort measurement invariance as described by Widaman, Ferrer, & Conger (2010), as shown in Table S1. There were no significant differences, by *χ*^2^ test, between models with configural, weak, strong, and strict measurement invariance. There were few differences in Akaike Information Criterion (AIC), while Bayesian Information Criterion (BIC) favoured the models with stricter factorial invariance. Thus, we considered the construct of g to be comparable across the cohorts, and proceeded with the mediation analyses.

We tested whether the paths from cohort to *g* (the “direct” path), from cohort to the mediator (the first part of the “indirect” path), and from the mediator to *g* (the second part of the “indirect” path) were significant, using 95% confidence intervals bootstrapped 1,000 times, to test whether the mediation (the product of the two “indirect” paths) was statistically significant. As an effect size, we calculated a “percentage of mediation” (Iacobucci et al., 2007), estimating how much the “direct” path was attenuated by the inclusion of the “indirect” path. We tested whether multiple mediators—for instance, grey matter, normal-appearing white matter, and white matter hyperintensity volumes—were incrementally significant mediators in a simultaneous model.

## Results

### Matching and health variables

We first matched the participants across the two cohorts using the propensity score procedure described above (the coefficients from the logistic regression used to generate the propensity score are provided in Table S2). To confirm the effectiveness of the propensity score matching procedure, we tested whether there were any significant differences in the matching variables. The results are shown in the upper section of Table 1. Differences in age-11 cognitive ability, background and own-attained social class and intracranial volume were small and non-significant (all *d*_*s*_ values < 0.17, all *p*-values > .36). The sex ratio did not differ between cohorts (*χ*^2^ = 0.00, *p* = 1.00). This confirmed that there were at most small, non-significant differences in these variables between the LBC1921 and LBC1936 participants.

**Table 1.** Comparisons between LBC1921 (*n* = 42; age ∼92) and propensity-score matched LBC1936 (*n* = 126; age ∼73) participants on each brain measure, for those who had data on all matching variables.

| Measure category | Measure | $n$ | | Sample mean (SD)/% | | Difference test | | |
| --- | --- | --- | --- | --- | --- | --- | --- | --- |
| | | LBC1921 | LBC1936 | LBC1921 | LBC1936 | $t$ | $p$ | $d_s$ |
| *Matching variables* | Father's SES | 42 | 126 | 2.90 (1.28) | 2.87 (1.00) | 0.15 | .88 | 0.03 |
|  | Achieved SES | 42 | 126 | 2.26 (0.99) | 2.41 (0.80) | −0.89 | .37 | 0.16 |
|  | Age 11 MHT | 42 | 126 | 47.36 (10.33) | 48.84 (12.45) | −0.76 | .45 | 0.14 |
|  | ICV | 42 | 126 | 1490.91 (140.18) | 1478.37 (141.80) | 0.50 | .62 | 0.09 |
|  | Sex | 42 | 126 | 40.5% male | 39.7% male | - | 1.00 | - |
| *Brain tissue measures* | TBV (cm<sup>3</sup>) | 38 | 126 | 954.73 (95.17) | 1015.49 (92.43) | −3.49 | <.001 | 0.62 |
|  | GMV (cm<sup>3</sup>) | 37 | 126 | 404.76 (39.02) | 480.20 (48.20) | −9.77 | <.001 | 1.74 |
|  | NAWM (cm<sup>3</sup>) | 37 | 126 | 434.02 (70.73) | 491.69 (54.05) | −4.58 | <.001 | 0.82 |
|  | WMH (cm<sup>3</sup>) | 40 | 126 | 46.87 (30.42) | 13.01 (12.01) | 6.81 | <.001 | 1.21 |
|  | MCT (mm) | 37 | 112 | 2.97 (0.14) | 3.13 (0.15) | −6.08 | <.001 | 1.08 |
|  | TSA (cm<sup>2</sup>) | 39 | 122 | 1377.11 (164.61) | 1572.46 (145.77) | −4.74 | <.001 | 1.12 |
| *Later-life cognitive tests* | Digit-Symbol | 34 | 125 | 33.32 (10.66) | 58.39 (11.75) | −11.89 | <.001 | 2.12 |
|  | Logical Memory | 42 | 126 | 9.90 (4.48) | 16.70 (3.77) | −8.85 | <.001 | 1.58 |
|  | Verbal Fluency | 42 | 126 | 38.43 (13.64) | 43.77 (12.00) | −2.26 | .02 | 0.40 |
| *Dementia screening* | MMSE | 42 | 125 | 26.90 (2.36) | 28.98 (1.29) | −5.45 | <.001 | 0.97 |

*Note:* Differences calculated using Welch's two-sample $t$ -test; for Cohen's $d_s$ , see Lakens (2013); LBC1921 = Lothian Birth Cohort 1921; LBC1936 = Lothian Birth Cohort 1936; SES = socioeconomic status; MHT = Moray House Test, ICV = intracranial volume, TBV = total brain volume, GMV = grey matter volume, NAWM = normal-appearing white matter volume, WMH = white matter hyperintensity volume, MCT = mean cortical thickness, TSA = total cortical surface area; MMSE = Mini-Mental State Examination.

Tables 2 and 3 compare both samples across multiple health variables. Compared to the matched younger LBC1936 sample, the older LBC1921 sample rated their health compared to one year ago more poorly (*χ*^2^ = 22.15, *p* < .001), and had higher rates of hypertension (76.2% vs. 42.1%, *χ*^2^ = 13.34, *p* < .001) and cancer/tumours (31.0% vs. 12.7%, *χ*^2^ = 7.35, *p* = .01). The older sample also had lower levels of blood haemoglobin (124.43g/L vs. 139.49g/L; *t* = –4.14, *p* < .001, *d*_*s*_ = 0.74) and eGFR (55.63ml/min vs. 63.84ml/min; *t* = –6.79, *p* < .001, *d*_*s*_ = 1.21). There were no significant differences between the cohorts in stroke prevalence (*p* = .13), or on any of the other health indicators (all *p*-values > .50). No participants in either cohort had a self-reported diagnosis of dementia.

**Table 2.** Comparisons between the LBC1921 (age ∼92) and the LBC1936 (age ∼73) on categorical health variables.

| Measure | Frequencies/Percentages | | $\chi^2$ | p-value |
| --- | --- | --- | --- | --- |
|  | LBC1921 (*n* = 42) | LBC1936 (*n* = 126) |  |  |
| Smoking status | Never smoked = 23<br>Ex-smoker = 16<br>Current smoker = 3 | Never smoked = 58<br>Ex-smoker = 59<br>Current smoker = 9 | - | .57 |
| Self-reported health | Poor = 2<br>Fair = 5<br>Good = 16<br>Very good = 16<br>Excellent = 3 | Poor = 1<br>Fair = 8<br>Good = 40<br>Very good = 62<br>Excellent = 15 | - | .19 |
| Self-reported health compared to 1y ago | Much worse than 1y ago = 4<br>Somewhat worse than 1y ago = 16<br>About the same as 1y ago = 22<br>Somewhat better than 1y ago = 0<br>Much better than 1y ago = 0 | Much worse than 1y ago = 0<br>Somewhat worse than 1y ago = 27<br>About the same as 1y ago = 81<br>Somewhat better than 1y ago = 13<br>Much better than 1y ago = 5 | - | <.001 |
| Hypertension | 76.2% | 42.1% | 13.34 | <.001 |
| Diabetes | 2.4% | 8.7% | 1.08 | .30 |
| Hypercholesterolemia | 38.1% | 43.7% | 0.20 | .65 |
| Cardiovascular disease | 50.0% | 33.3% | 3.06 | .08 |
| Leg pain | 54.8% | 40.5% | 2.06 | .15 |
| Blood circulation problems | 16.7% | 22.2% | 0.30 | .58 |
| Stroke | 11.9% | 7.1% | 4.03 | .13 |
| Cancer/tumour | 31.0% | 12.7% | 7.35 | .01 |
| Thyroid disorder | 14.3% | 12.7% | 0.00 | 1.00 |
| Parkinson's disease | 0.0% | 0.8% | 0.00 | 1.00 |
| Dementia | 0.0% | 0.0% | 0.00 | 1.00 |
| Arthritis | 42.9% | 42.9% | 0.00 | 1.00 |

Note: $p$ -values from Fisher's Exact Test (initial 3 rows), or from chi-squared test with Yates's continuity correction (remaining rows). One LBC1921 participant who was unsure if they had been diagnosed with stroke was classified as not having such a diagnosis for this table.

**Table 3.** Comparisons between the LBC1921(age ∼92) and the LBC1936 (age ∼73) on continuous health (biomarker) measures.

| Measure | Sample mean (SD) |  | Difference test |  |  |
| --- | --- | --- | --- | --- | --- |
|  | LBC1921 (*n* = 42) | LBC1936 (*n* = 126) | *t* | *p* | *d<sub>s</sub>* |
| Blood haemoglobin | 124.43 (11.95) | 139.49 (13.48) | -6.79 | <.001 | 1.21 |
| White cell count | 7.01 (2.05) | 6.60 (1.47) | 1.18 | .24 | 0.21 |
| Fibrinogen | 3.16 (0.76) | 3.33 (0.66) | -1.24 | .22 | 0.22 |
| D-dimer | 376.91 (462.24) | 206.71 (197.71) | 1.98 | .055 | 0.35 |
| eGFR | 55.63 (12.33) | 63.84 (3.93) | -4.14 | <.001 | 0.74 |
| HbA1c | 5.61 (0.40) | 5.74 (0.59) | -1.58 | .12 | 0.28 |

Note: Differences calculated using Welch's two-sample *t*-test; *d*-values are Cohen's *d<sub>s</sub>* (see Lakens, 2013); eGFR = estimated Glomerular Filtration Rate; HbA1C = glycated haemoglobin.

### Brain measure comparisons

There were substantial differences between the two cohorts in brain volume, grey matter volume, normal-appearing white matter volume, white matter hyperintensity volume, mean cortical thickness, and total cortical surface area (Table 1). The age-92 LBC1921 had significantly and substantially lower volumes than the age-73 LBC1936 on all healthy brain tissue measures (all *p*-values < .001), with effect sizes (Cohen’s *d*_*s*_) between 0.62 and 1.74. The older LBC1921 participants had significantly and substantially higher volumes of white matter hyperintensities (*p* < .001, *d*_*s*_ = 1.21). The standardized differences are illustrated in Figure 1.

**Figure 1.**
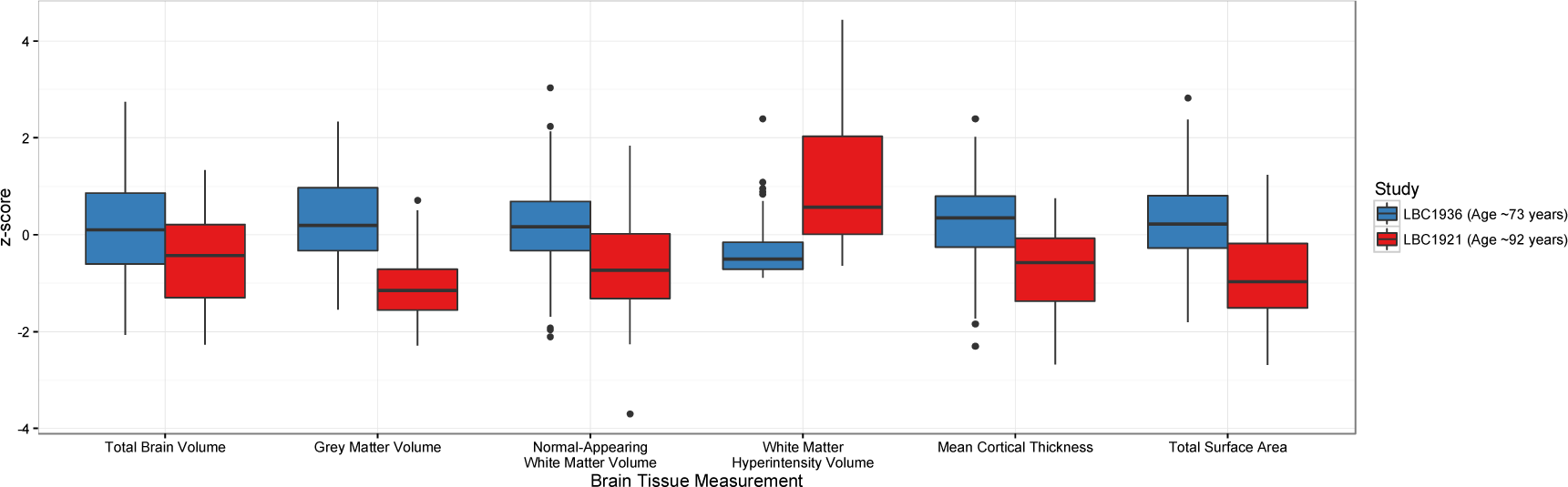
Brain tissue differences between the LBC1936 (age ∼73) and the LBC1921 (age ∼92) for each tissue type, on a standardized (*z*) scale. *p*-values for all comparisons are < 0.001 (see Table 1).

### Subregional brain differences

We next compared brain measures from the two cohorts at the level of subregions. We examined three different measures: subregional volume, subregional surface area, and vertex-wise cortical thickness.

#### Cortical volume and surface area

The older cohort exhibited significantly lower volumes and smaller surface areas across almost all cortical regions (see Figure 2, and Tables S2 and S3). The only regions that showed no detectable differences were the volumes of the frontal poles, the temporal poles and portions of the bilateral cingulate and right paracentral cortex, and the surface areas of the left frontal pole and the right insula and paracentral cortex. The most pronounced effect was found in the bilateral temporal lobes on both the lateral surface (volume: all *p*-values < .001, all *d*_*s*_ values > 0.85; and surface area: all *p*-values < .001, all *d*_*s*_ values > 0.83) and medial surface (volume: all *p*-values < .001, all *d*_*s*_ values > 0.73; and surface area: all *p*-values < .001, all *d*_*s*_ values > 0.97). The left inferior temporal cortex showed the largest effect sizes of all (volume: *p* < .001, *d*_*s*_ = 1.63; and surface area: *p* < .001, *d*_*s*_ = 1.53). There were also strong group differences in the surface area of the frontal lobes (all *p*-values < .001, all *d*_*s*_ values > 0.97). Significant group effects were generally of lowest magnitude for somatosensory, motor and cingulate regions.

**Figure 2.**
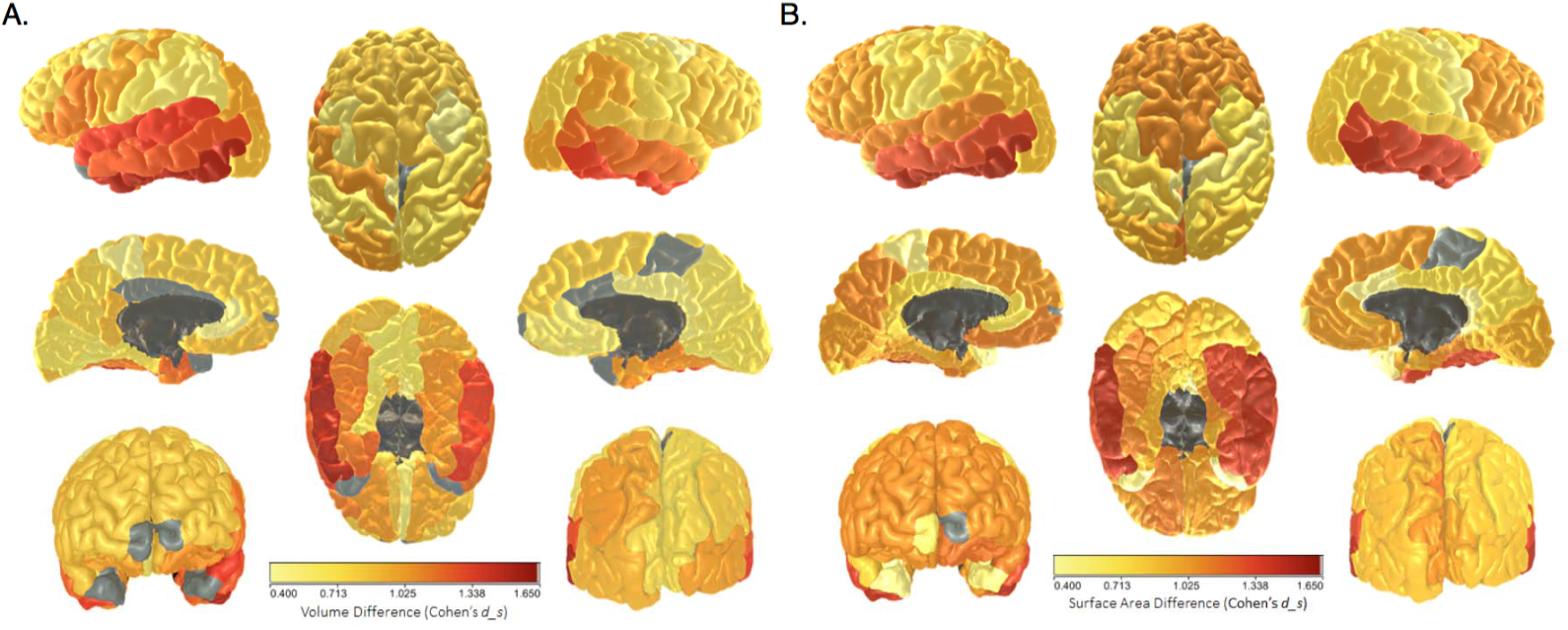
Differences between the LBC1936 (age ∼73) and the LBC1921 (age ∼92) in A. mean volume and B. mean surface area across parcellated subregions. Grey indicates null difference. For full numerical results for volume and surface area respectively, see Tables S2 and S3. Light grey denotes non-significant associations, dark grey denotes unlabelled regions.

#### Cortical thickness

There were vertex-wise differences between the cohorts across most of the cortical mantle, with the strongest differences localized to the superior temporal lobe/insular cortex (Figure 3).

**Figure 3.**
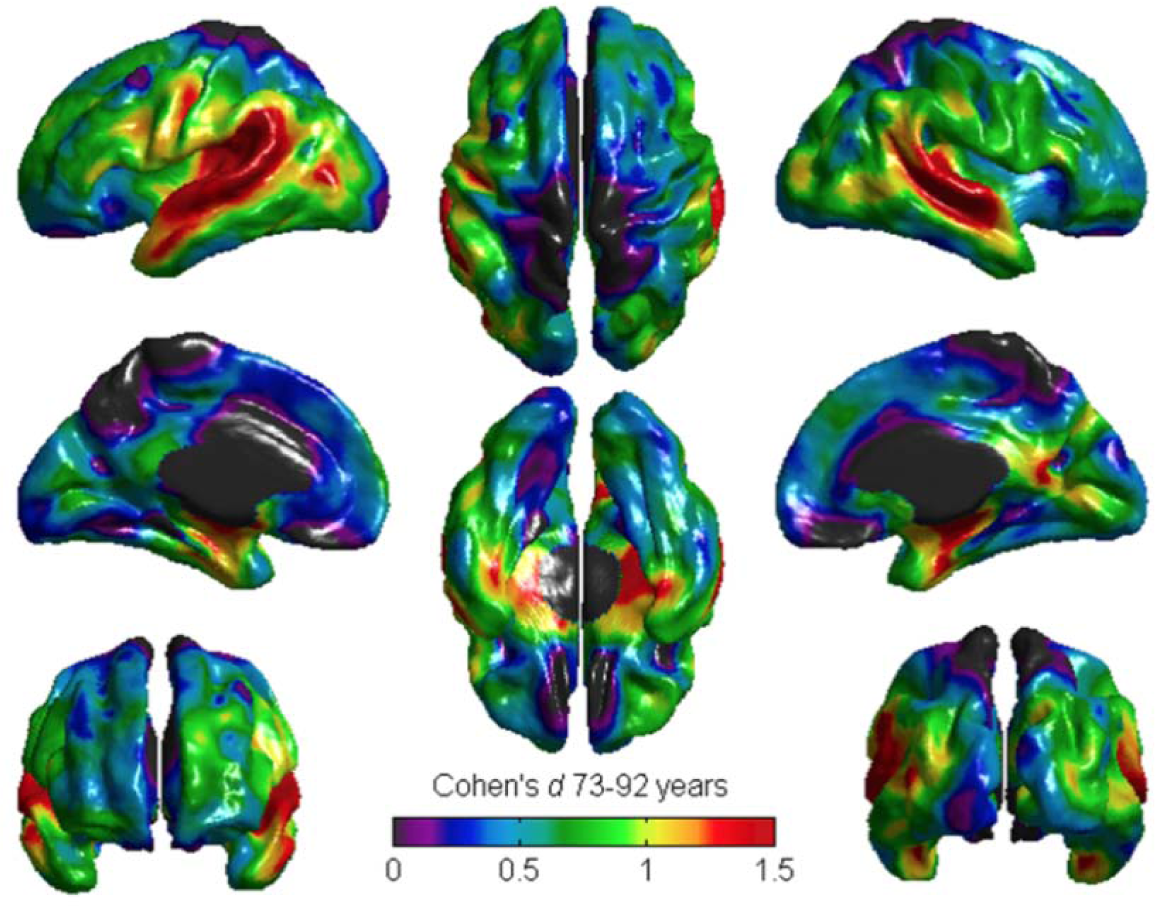
Vertex-wise differences in cortical thickness between the LBC1936 (age ∼73) and the LBC1921 (age ∼92). Warm and grey colored areas indicate that the greatest standardized differences between groups (*d* > 1.5 SD) were localized to the superior temporal lobe. Black denotes unlabelled regions.

#### White matter hyperintensity location

Probability distribution maps of WMH comparing the two cohorts showed that the WMH from the younger sample (LBC1936; age 73) were mainly concentrated in the periventricular regions, particularly at the horns of the lateral ventricles. In the older sample (LBC1921, age 92), WMH were additionally found in the deep white matter regions and were particularly abundant in the centrum semiovale. These differences are illustrated in Figure 4.

**Figure 4.**
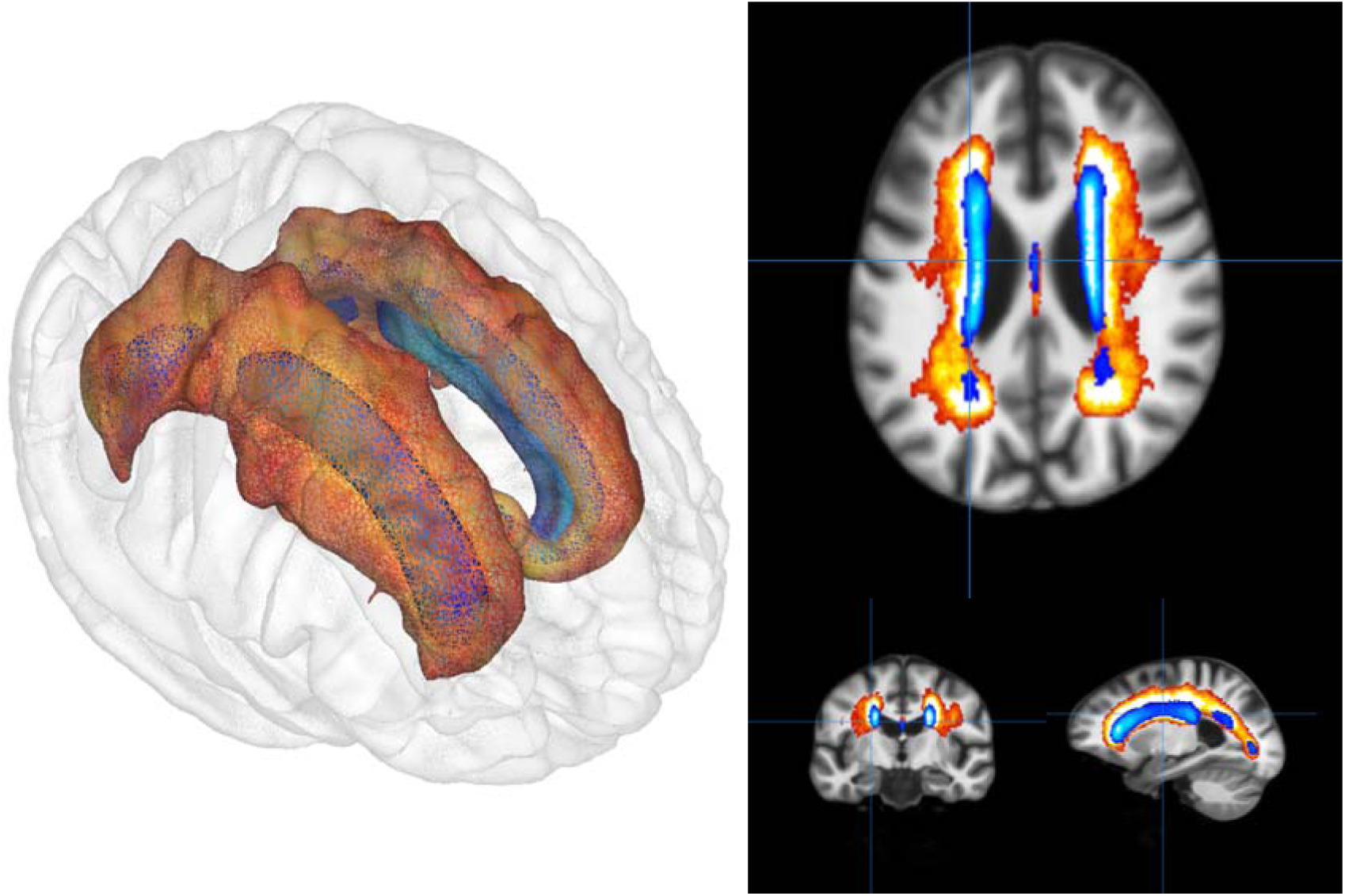
Comparison of the white matter hyperintensity (WMH) probability maps for the age-73 LBC1936 group (blue) and age-92 LBC1921 group (red). Shown are axial, coronal, and sagittal views, along with a 3D render of the locations. Lighter colours within each cohort indicate a greater probability of WMH being found in that location.

### Cognitive mediation models

There were substantial and significant cognitive differences between the two cohorts in later life (lower part of Table 1). The age-92 LBC1921 performed more poorly than the age-73 LBC1936 on all measures, particularly so for the Digit-Symbol Substitution and Logical Memory tests (*d*_*s*_-values = 2.12 and 1.58, respectively), and less so for the Verbal Fluency test (*d*_*s*_ = 0.40). There was also a substantial average difference in MMSE scores: (*d*_*s*_ = 0.97), though only four participants, all of them in the LBC1921, had scores below the cutoff of 24 (two with a score of 23, one with a score of 22, and one with a score of 21).

We tested the extent to which brain-level differences mediated the cohort (that is, age) differences in a general cognitive ability (*g*) factor formed from performance on the three normal-range cognitive tests (Digit-Symbol Substitution, Logical Memory, and Verbal Fluency; see Table S5 for a correlation matrix showing each of the brain and cognitive variables within each cohort). We first tested each of the brain variables as separate mediators. The coefficients from each of these models, all of which had adequate or good fit to the data (all Root Mean Square Error of Approximation (RMSEA) values < 0.094, all Comparative Fit Indices (CFI) > 0.974; all Tucker-Lewis Indices (TLI) > 0.936). Individually, the only variables that did not significantly mediate the relation between cohort and *g* were total brain volume and mean cortical thickness (for all other variables, the bootstrapped 95%CIs did not include zero; Table S6). We then tested models including multiple previously-significant mediators to investigate which variables mediate additional variance over and above one another. In models that also included white matter hyperintensity volume, the mediation paths via grey matter and normal-appearing white matter volumes were reduced to non-significance (*β*_mediation_ [95%CIs]: .05 [-.07, .16] and .01 [-.06, .07], respectively). However, this was not the case for total surface area: as shown in Figure 5, both white matter hyperintensity volume (*β*_mediation_ = .13 [.05, .23]) and total surface area (*β*_mediation_ = .07 [.003, .13]) were independently significant mediators (with the opposite direction of effects: as expected, lower white matter hyperintensity volume and greater surface area were related to higher *g*). Together, the two brain structural measurements of total surface area and volume of white matter hyperintensities mediated 24.9% of the relation between cohort (age) and *g*.

**Figure 5.**
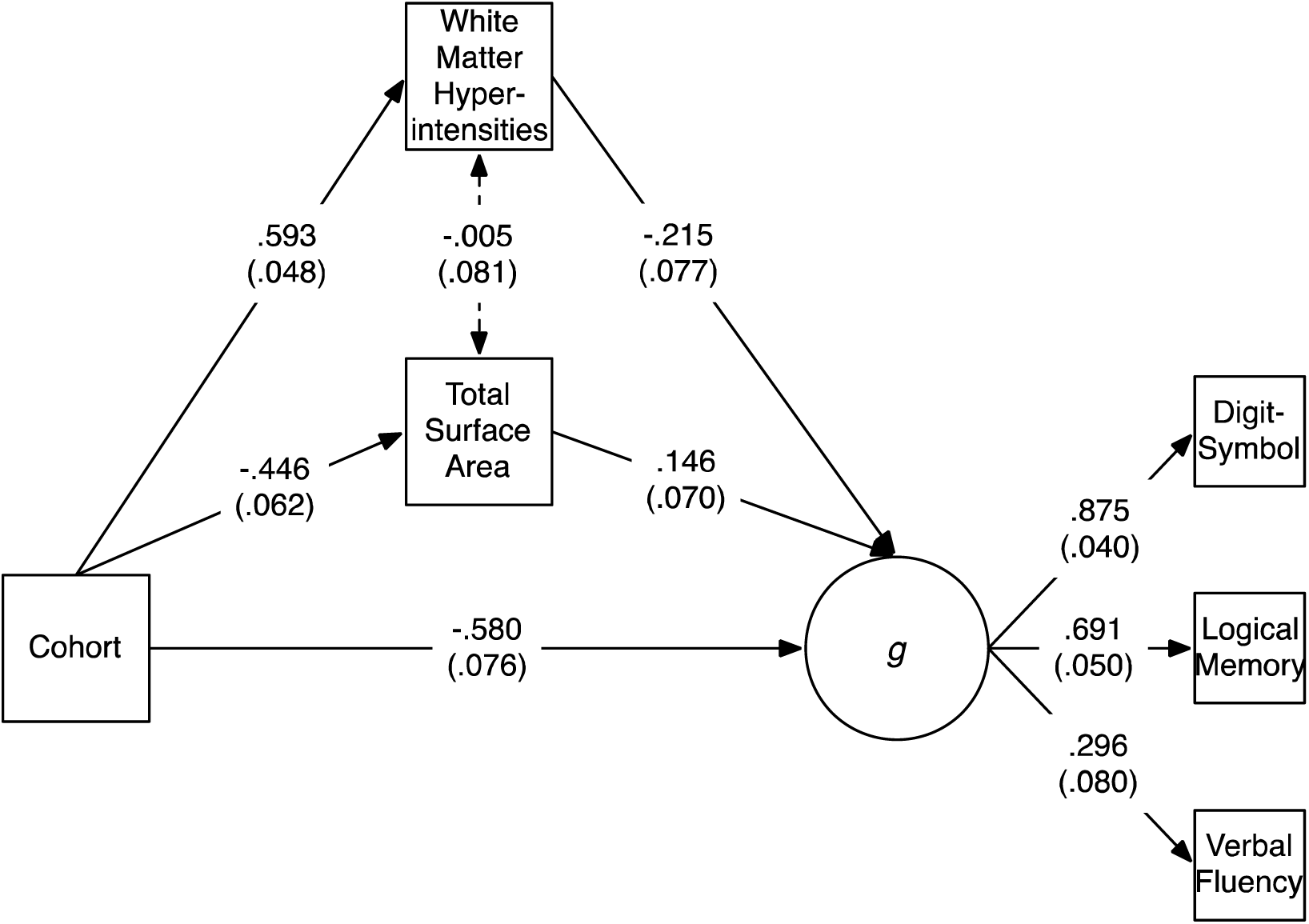
Mediation model including white matter hyperintensity volume and total surface area as mediators between cohort (age) and *g* (general cognitive ability, as indicated by three cognitive tests). Values are standardized path coefficients with standard errors in parentheses. The dotted line between the two neural variables indicates a statistically non-significant relation.

For comparison, we ran the same mediation analyses after choosing not a propensity-score-matched comparison sample of LBC1936 participants, but a sample chosen entirely at random from the full cohort (we thank an anonymous reviewer for suggesting this analysis). For every mediator except WMH, the mediator explained a smaller percentage of the cohort-*g* relation. This indicates that, overall, the age-*g* association is somewhat more strongly mediated when the age groups are more similar in their background characteristics, possibly because there is less noise in the estimation of the cohort-brain relation.

In addition, we ran an analysis where we controlled the multiple-mediation model (including WMH and total surface area as moderators) for vascular risk factors. This was carried to test whether the mediation by WMH was attenuated after control for these factors (WMH are hypothesized to be vascular-related). We created a sum score of the following self-reports: smoking (1 point for being an ex-smoker, 2 points for being a current smoker), hypertension, diabetes, hypercholesterolemia, cardiovascular disease, and stroke. This ‘vascular risk score’ ran from 0 to 6—no participants had the maximum score of 7—with an average of 2.05 (SD = 1.33). It had a zero-order correlation of *r*(164) = .17, *p* = .03 with WMH volume. We then re-ran the mediation model including WMH and total surface area, regressing both mediators and the outcome on the vascular risk score. This made essentially no difference to the mediation: all paths were still significant with barely-changed effect sizes, and the overall percentage mediation of the cohort-*g* association from WMH and total surface area was 26.0%.

### Analyses adjusting for the Flynn Effect

In a comparison across the two Scottish Mental Surveys, it was noted that the Moray House Test score of all Scottish 11-year-olds in 1947 was on average 2.284 points higher than the score of all Scottish 11-year-olds in 1932 (Deary, Whalley, & Starr, 2009). This reflects the well-known ‘Flynn Effect’, which is the tendency for IQ test scores to increase generation-upon-generation (e.g. Flynn, 1987; Pietschnig & Voracek, 2015). Since the MHT score was used as a matching variable, the Flynn Effect may have affected our cohort matching. For this reason, we re-ran the comparisons shown in Table 1 after adding 2.284 points to the MHT score of all the participants in the older (LBC1921) sample. This resulted in a slightly different sample of matched individuals from the LBC1936. The results are shown in Table S7 in the Supplementary Materials: they are near-identical to the results from the unadjusted analysis. Probably because other variables were used for matching in addition to the MHT scores, the Flynn Effect appears not to have made a substantial difference to the comparisons in this study.

## Discussion

This propensity-score matching study compared 92-year-olds with a matched group of 73-year-olds on a variety of structural brain measures. The 19-year difference in their ages was linked to substantial differences in brain structure. The older participants had less healthy brain tissue and higher volumes of white matter hyperintensities (Figure 1), which spread over a substantially larger area (Figure 4). Their mean brain cortical thickness and total cortical surface area were lower. These differences were large in size, often upwards of a full standard deviation (for example, grey matter volume was 1.74 SDs lower in the older group), and the specific differences in white matter hyperintensities and cortical surface area mediated around a quarter of the association between age and general cognitive ability. The findings illustrate, using a valuable and rarely-available study design, tissue- and region-general associations of age with brain structure.

Comparing our three subregional analyses—of subregional volumes, subregional surface areas, and vertex-wise cortical thickness—reveals broadly similar patterns of age differences. For instance, differences in frontal areas were strongly related to age, but we found that the temporal lobes showed even larger age associations. The temporal lobe is often highlighted as an area of particular importance for Alzheimer’s Disease (AD), with many studies noting that AD affects the temporal lobe much more prominently than normal ageing (e.g. Bakkour et al., 2013; Dickerson et al., 2009; Fox & Schott, 2004; Fjell & Walhovd, 2010). The effect sizes observed here (for example, a negative difference of *d*_*s*_ = 1.63 for the inferior temporal area across 19 years; Table S3) may serve as a baseline for future studies and comparative diagnoses; although none of our participants had a diagnosis of AD, they still showed strikingly large differences in their temporal lobes, raising the possibility that differences in this area may be a less useful indicator of AD (though see below for discussion of the results of the dementia screening test).

There were substantial differences in cognitive abilities between the samples: these were less prominent for verbal fluency, but were very strong for logical memory, a measure of verbal declarative memory, and even more so for digit-symbol substitution, a measure of cognitive processing speed (Salthouse, 1996). These age differences were reflected in a large association of cohort (in this study, a strong proxy for age) with the latent *g*-factor of cognitive ability in the mediation model (Figure 4). The healthy brain tissue volumes no longer mediated significant amounts of variance after the volume of unhealthy tissue (white matter hyperintensities) was included. In a review of age-brain-cognitive relations that included 254 results from mediation analyses, Salthouse (2011) noted that there was little replicable evidence for healthy tissue volume’s mediation of age-cognitive relations; our results would appear to be consistent with this conclusion, though this only became clear after the variable of white matter hyperintensity volume was added to the model.

As we have found in previous longitudinal research (Ritchie et al., 2015a), the extent of white matter hyperintensities, which our lesion-mapping analysis also showed to be far larger in the older participants, appears to be the best brain structural indicator of cognitive health, at least of those considered here. Again, our results are consistent with the conclusion drawn in the review of Salthouse (2011), who argued that white matter hyperintensities mediated the age-related variance in many different cognitive tests (see also Rabbitt et al., 2007); on the basis of our results, where white matter hyperintensities significantly mediated the age relation with general cognitive ability (*g*), we would tend to agree; however, we were unable to go further and examine specific domains of cognitive ability because we only had available three cognitive tests that were taken by participants in both cohorts. Consistent with some of our previous research in the LBC1936 cohort (Dickie et al. 2016b; Wardlaw et al., 2014), the vascular risk factors measured here, such as hypertension, smoking, diabetes, and hypercholesterolemia, accounted for only a very small proportion of the variance in brain and cognitive variables, implying that we may have to look elsewhere for health and lifestyle predictors of brain and cognitive differences in older age.

One non-volumetric measure independently mediated age variance in the *g*-factor in addition to white matter hyperintensities: total cortical surface area. Cortical thickness, on the other hand, was no longer significant in the mediation model after inclusion of white matter hyperintensities. This is despite it making incremental contributions to explaining *g* level beyond hyperintensities in a previous paper which used the full sample of LBC1936 participants, all at age 73 (Ritchie et al., 2015b). There was, however, good reason to predict that cortical thickness and surface area would have separable effects: not only were they weakly and non-significantly correlated in both the present cohorts (see Table S5), but previous longitudinal research has found them to age on distinct trajectories (Hogstrom et al., 2013; Storsve et al., 2014). Genetically-informative studies have also found them to be under dissociable genetic influence (Panizzon et al., 2009; see also Eyler et al., 2012; Winkler et al., 2010). Surface area and cortical thickness have also been shown to have different relations with intelligence during development (Schnack et al., 2015).

In their discussion of this issue, Fjell et al. (2014) note that we currently have only a poor understanding of the reasons underlying this apparent dissociation of surface area and cortical thickness (though see Seldon, 2005, for one speculative mechanism regarding the ‘stretching’ of the cortex during its development).

Overall, the effect size of the mediation by white matter hyperintensities and surface area (26.4% of the cohort-*g* association) was substantial, but it leaves a large portion of the relation between age and cognitive ability unaccounted-for. Different or more detailed brain measurements, in larger samples so as to increase statistical power to detect smaller significant contributions, are required. Measures of the brain’s white matter microstructure, whether in terms of tractography (e.g. Clayden et al., 2011; which were taken in both cohorts here but were not comparable due to the lower-resolution diffusion data obtained for the older cohort) or connectomics (which can also be examined using functional imaging; Bullmore & Bassett, 2010), as well as newer measures such as cortical complexity (Madan & Kensinger, 2016), are likely to have a role to play in explaining further portions of the age-cognitive relation. It would also be of interest, in a larger sample, to extend our mediation analyses, which only used broad measures from across the brain, to the subregional context, investigating which brain subregions (and which measures of them) mediate the largest proportion of the relation between age and cognitive ability.

### Strengths and limitations

The strengths of the study include the availability of our two narrow-age samples without (self-reported) dementia, one of which was in the tenth decade of life. Importantly, both samples had data on the same test of general intelligence, taken at the same age in childhood. All of the later-life cognitive and health measures were also taken comparably, and the same MRI scanner was used for all participants’ scans, albeit with small differences in image resolution arising from the necessity to provide an imaging protocol of ∼30 minutes suitable for scanning participants in their nineties (LBC1921). The propensity score matching technique allowed us to select three similar participants from the larger LBC1936 sample for each LBC1921 participant.

The present study also includes a number of limitations that may produce biases in the results and affect their generalizability to other populations of older individuals. One important limitation of the study is the sample size of the older (LBC1921) group: we may not have had sufficient power to detect significant differences in the matching variables (though the absolute effect sizes of these differences were all relatively small). The small sample will have reduced the precision of our estimates. However, as noted above, it is rare to have a sample of individuals with MRI data who have all reached the age of 92 years, and they were each matched with three members of the younger, age-73 cohort. The small sample also prevented us from examining differences in the distributions of the variables as well as in the means: future research with larger samples should test for variance differences in older participants, as was recently found for white matter diffusion measures within a large cross-sectional sample (e.g. Cox et al., 2016).

Sample selectivity is also an important issue: especially in the older sample, the fact that they attended for testing and MRI scanning itself implies that the participants were likely to be healthier than the average member of the general older-age population. This means that we will not have included individuals with more severe illness and frailty, and potentially have underestimated the effect sizes we report here. Relatedly, more individuals in the LBC1921 group showed possible mild cognitive impairment or dementia according to the Mini-Mental State Examination, despite not reporting having received a formal diagnosis of dementia. Although the number of participants who were below the cutoff (four of the total forty-two) was small, and none were substantially below the cutoff (lowest score = 21) it should be taken as an indication that ageing-related pathologies may have been more commonly present in the older sample, potentially reducing the proportion of the neuroanatomical and cognitive differences found here that were entirely due to the normal ageing process.

A further limitation concerns cohort effects. As we have previously noted (Deary & Ritchie, 2016), the older cohort were adults by the time of the Second World War, and experienced fewer years of life with free healthcare available on the United Kingdom’s National Health Service (which was founded in 1948). The potential effects of these differences, combined with those of a large number of other social, technological, and medical advances across the 20^th^ Century, are near-impossible to control for. Naturally, all studies comparing older and younger groups suffer from similar limitations, though we were fortunate in having high-quality data on how the participants performed cognitively in childhood to provide a baseline for comparison.

We used a validated segmentation procedure and individually checked all tissue masks for each subject in both cohorts to minimize the effect of measurement error. However, some degree of error (for instance, due to motion, registration, or image resolution), will always be present in quantitative measurements of brain tissues. Finally, we did not quantitatively measure skull thickening in these groups, or control for its influence: this is likely to occur between ages 73 and 92 (Aribisala et al., 2014), meaning our use of intracranial volume as a proxy for maximal brain volume is less reliable in the older sample.

### Conclusion

Cross-sectional comparisons of ageing are often limited by the lack of knowledge of how similar the participants were earlier in life. Here, we were able partly to avoid this limitation by matching our participants on several background characteristics, including childhood cognitive ability (a valuable predictor of various important late life outcomes). Examining differences in several brain measures between matched narrow-age groups aged 73 and 92 years, we observed neurostructural differences that were often very large in size. We found that two of the neuroanatomical measures, white matter hyperintensities and total cortical surface area, mediated one quarter of the relation between age and general cognitive ability. Although the older sample in particular was small, our results provide a measure of what can be expected neurostructurally across this period of life from a unique perspective, and raise the question of which other neuroimaging measures might more fully mediate the negative relation between age and cognitive ability.

## Acknowledgements

We thank the Lothian Birth Cohort 1921 and 1936 participants and research teams. We are grateful to the Scottish Council for Research in Education for access to the data from the Scottish Mental Surveys of 1932 and 1947. We thank the nurses, radiographers, and other staff at the Wellcome Trust Clinical Research Facility and the Brain Research Imaging Centre in Edinburgh. The Lothian Birth Cohort 1921 data collection was supported by the Biotechnology and Biological Sciences Research Council (BBSRC; 15/SAG09977) and by a Royal Society-Wolfson Award to author I.J.D. The Lothian Birth Cohort 1936 data collection was supported by Age UK (Disconnected Mind project). The work was undertaken by The University of Edinburgh Centre for Cognitive Ageing and Cognitive Epidemiology, part of the cross-council Lifelong Health and Wellbeing Initiative (MR/K026992/1). Brain imaging analysis was supported by the Medical Research Council (MRC; G1001245 and G0701120), which also supports author S.R.C. (MR/M013111/1). Author J.M.W. is partly funded by the Scottish Funding Council as part of the SINAPSE collaboration. We thank Dr. Elliot Tucker-Drob for suggesting the analyses adjusting for the Flynn Effect.

